# Collagen Deposition in Diabetic Kidney Disease Boosts Intercellular Signaling: A Mathematical Model

**DOI:** 10.1101/2021.03.25.437068

**Authors:** Haryana Y. Thomas, Ashlee N. Ford Versypt

## Abstract

Diabetic kidney disease is a health burden that is becoming more prevalent in the US and worldwide. The limited options for treating and preventing diabetic kidney disease are in part due to gaps in our understanding of the progression of diabetic kidney damage and its impacts on cellular function. An important cellular function in the kidney glomerulus is intercellular communication via the release and uptake of soluble cytokines and growth factors. In diabetic kidney disease, excess collagen deposition alters the mesangial matrix properties, which, we hypothesize, diminishes the intercellular signaling between key glomerular cells. To test our hypothesis, we utilized established mathematical models of transport to study the impact of pathological deposition on the ability of cells to communicate via intercellular signaling. Our analysis reveals that pathological collagen deposition can enhance the signaling range of the glomerular cells rather than diminishing it.

**SIGNIFICANCE:** The incidence of diabetes is expected to rise to over 600 million by the year 2040, one third of whom are expected to develop diabetic kidney disease. Our lack of understanding of how diabetic kidney disease progresses and affects cellular and tissue function has led to our inability to mitigate the rapidly rising burden of diabetic kidney disease. A hallmark of diabetic kidney damage is collagen deposition, yet its impact on cellular and tissue function is still not well understood. The elucidation of the impact of collagen deposition on cellular and tissue function will enable the identification of mechanisms that exacerbate the progression of diabetic kidney damage and thus provide novel avenues for preventing or slowing the progression of diabetic kidney damage.

## INTRODUCTION

Diabetes is a significant burden on public health. In 2015 over 400 million people had been diagnosed with diabetes worldwide, a number that is expected to rise to over 600 million by 2040 (1). One third of these diabetic patients are expected to develop diabetic kidney disease, the leading cause of kidney failure (1, 2). Our lack of understanding of how diabetic kidney disease progresses and affects cellular and tissue function has contributed to our inability to mitigate the rapidly rising burden of diabetic kidney disease. Characterized by proteinuria and a loss in glomerular filtration rate, diabetic kidney disease is induced by biochemical perturbations and morphological changes such as excess collagen deposition in the kidney mesangium (3, 4). Although excess collagen deposition in the mesangium is a hallmark of diabetic kidney damage, its impact on cellular and tissue function is still not well understood. Thus, our aim is to use a modeling approach to fill this gap in knowledge.

The mesangium is a collagenous matrix that is located at the center of the glomerulus, the filtration unit of the kidney. Centrally located, the mesangium is the region that separates the three key glomerular cell types: mesangial, endothelial and podocyte cells. These cells are in constant communication in health and disease (2, 5). Studies have shown that signaling between endothelial and mesangial cells is crucial for the formation of a healthy glomerulus. Elimination of PDGF-*β*, a key signaling molecule between mesangial and endothelial cells, has been shown to cause the development of an abnormal glomerulus (6). This finding has been corroborated by multiple studies (7–9). Additionally, inhibition of signaling pathways that mediate crosstalk between glomerular cells have been shown to ameliorate glomerular diseases (10, 11). Consequently, we can see that interference in the signaling between glomerular cells has an impact on the function of the glomerulus in both health and disease. Interference in glomerular cell signaling occurs during diabetic kidney damage when the ability of signaling molecules to traverse through the mesangium is altered by excess collagen deposited in the mesangium. The extent to which the collagen deposition impacts intercellular signaling between glomerular cells has not been previously investigated. We hypothesize that the pathological deposition of collagen blocks the path of signaling molecules and thus decreases the ability of glomerular cells to communicate.

Previously, mathematical models have been developed to study cell-cell communication. Francis et al. developed a mathematical model to study the effective signaling distance of a single, hybridoma cell (12, 13). They found that a hybridoma cell can effectively signal on the order of 250 *μ*m within a time span of 10-30 mins. They also used their mathematical model to show that the effective signaling distance is governed by a key dimensionless group consisting of the diffusivity of the signaling molecule, the secretion rate, and the binding constant of the signaling molecule to a receiving cell (12). The model by Francis et al. (12) has been extensively modified to study the impact of varying constant and pulsatile secretion rates and signaling molecule degradation on the effective signaling distance (14–18). In this work, we utilize the model of Francis et al. (12) to study the impact of pathological collagen deposition on the effective signaling distance of a mesangial cell. In addition to Francis et al.’s modeling approach, we use an empirical relationship (19) to quantify the changes in diffusivity of signaling molecules that occurs due to the pathological collagen deposition. We find from our quantitative study the surprising result that pathological collagen deposition can actually enhance the ability of cells to communicate rather than diminishing their ability to communicate.

## METHODS

### Solitary Cell Model

In this model, we consider a single mesangial cell secreting signaling molecules that diffuse radially outward from the cell surface into the extracellular matrix. A one dimensional, spherical mass balance equation

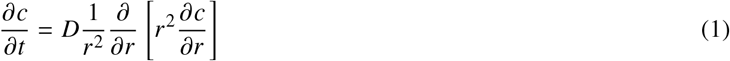

captures the temporal and spatial dynamics of the diffusing signaling molecules, where *c* represents the concentration of the signaling molecules, *D* represents the diffusion coefficient for signaling molecules diffusing through the matrix, *r* represents the radial distance from the center of the cell, and *t* represents time (12). The cell is assumed to be a spherical object of radius *ρ* residing in an isotropic matrix. We recognize that *D* will change over the course of mesangium fibrosis in diabetic kidney disease, but it is assumed to be spatially uniform at various snapshots along the trajectory of disease progression, represented by different *D* values used in this study.

The distribution of the signaling molecules through the matrix is dependent on physiologically relevant initial conditions and boundary conditions. These initial and boundary conditions are summarized below and are teh same as those applied by Francis et al. (12). The first boundary condition

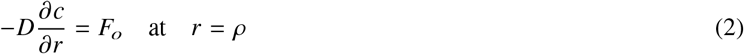

is a constant secretion rate, *F*_*o*_, at the cell surface. The second boundary condition is

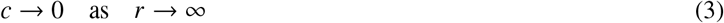

We specify that initially there are no signaling molecules present in the matrix

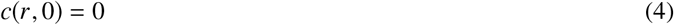

Given these conditions, an analytical solution that describes the concentration profile of the signaling molecules in time and space can be obtained (12, 18):

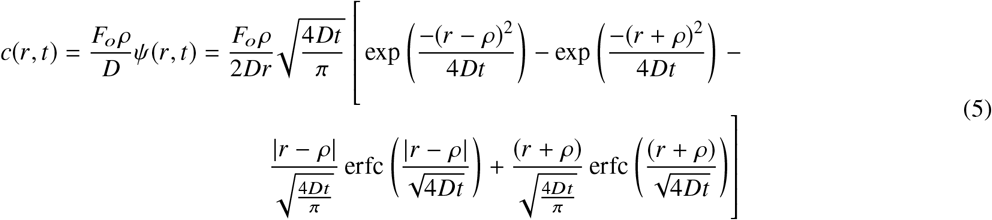

The key parameters in Eq. 5 are *F*_*o*_, *D*, and *ρ*.

### The Effective Signaling Distance

The effective signaling distance is obtained by finding the steady state propagation distance of the threshold concentration *k*_*m*_, which is the minimum concentration required to induce a response in the receiving cell. The propagation of the threshold concentration over time can be tracked by solving

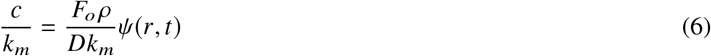

for *r* (12). The steady state distance propagated by the threshold concentration can be found by taking the limit of Eq. 6 as time approaches infinity (12)

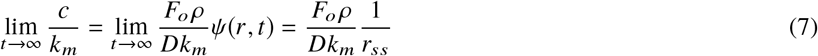

The steady state distance *r*_*ss*_ when *c* = *k*_*m*_ is

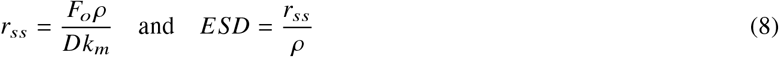

which we define to be the effective signaling distance (ESD). Francis et al. defined the effective signaling distance to be half the maximum propagation distance because it took over 10^8^ seconds to reach steady state in their scenario, which is an unrealistically long time scale for intercellular communication (12). In contrast, we define the effective signaling distance at the steady state distance because this steady state value is attained much faster for our case due to the much smaller secretion rate of mesangial cells in comparison to hybridoma cells. Since the effective signaling distance is dependent on diffusivity, as shown in Eq. 8, we use it to study how changes in diffusivity of signaling molecules through the remodeling matrix impacts the effective signaling distance.

### Diffusion Through a Remodeling Matrix

The diffusivity of signaling molecules is dependent on the properties of the matrix and the signaling molecules. In this section we use a model of diffusion through polymer hydrogels composed of rigid fibers (19) to model the diffusivity of signaling molecules through the mesangial matrix. Since the mesangial matrix is composed of rigid collagen fibers, the model of diffusion through polymer hydrogels can be used to approximate the diffusion of signaling molecules through the mesangial matrix. The model takes into account the hydrodynamic drag and obstruction effects of matrix fibers on solute molecule diffusivity (19) and is a model that has been previously used to capture the dynamics of solute diffusivity through the extracellular matrix of the mesangium (20). The matrix is modeled as a fibrous mesh with a specific, physiologically relevant porosity and average fiber radius. Collagen deposition is assumed to change the porosity of the matrix, which in turn changes the effective diffusion coefficient *D* via the relationship (19)

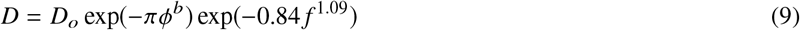

*D*_*o*_ is the free diffusion coefficient given by the Stokes-Einstein equation

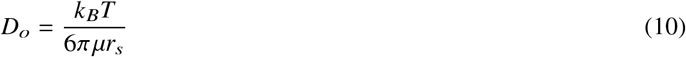

where *k*_*B*_ is Boltzmann’s constant, *T* is the temperature, and *μ* is the viscosity of interstitial fluid, which we assume to have the same viscosity as blood plasma. The following functions define the terms *b* and *f* in Eq. 9:

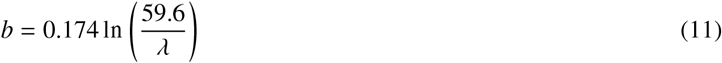

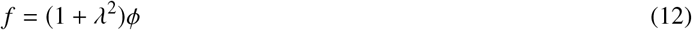

where *λ* is the the ratio of the solute radius, *r*_*s*_, to the fiber radius, *r*_*f*_.

### Parameter Selection

Model parameters were obtained from previous papers that modeled transport through the mesangial matrix. In cases where exact parameters values were unavailable, we found general expected ranges. For instance, mesangial cell cytokine secretion rates are not readily available in the literature because quantification of secretion rates is difficult (21, 22) and the methods are still being refined (21, 23–26). For short range signaling cells, however, cytokine secretion rates are estimated to be on the order of 10-100 molecules/cell/s (25, 27, 28). Threshold concentrations also vary widely but are estimated to be on the order of 10-50 pM (12). We chose the upper end of the secretion rates and lower end of the threshold concentration to obtain the upper limit of the signaling ranges. Collagen volume fractions in kidney fibrosis as well as fibrosis in other tissues is on the order of 0.05 to 0.30 (29–31). A majority of cytokines and growth factors have molecular weights that are on the order of 10-60 kDa (32, 33), which approximates to hydrodynamic radii of 2-6 nm.

### Numerical Methods and Code Repository

The concentration profile Eq. 5, propagation distance Eq. 6, and effective diffusivity calculations Eq. 9–Eq. 10 were solved using MATLAB version R2020b. The MATLAB code used for the calculations has been made available in an online repository at github.com/ashleefv/ColDepSignalingImpact (35).

## RESULTS AND DISCUSSION

### Approaching the Steady State Distance

Using the parameters listed in Table 1, we obtained a concentration profile of the signaling molecule through the matrix at specific times for 10^8^ seconds, a period long enough for steady state to be achieved (Figure 1a). Over time, as expected, signaling molecules begin to accumulate as the cell continues to secrete. More importantly, as the time progresses the accumulation rate of signaling molecules decreases and approaches a steady state value of around two and a half times the threshold concentration at the cell surface (Figure 1a). This approach to steady state is more clearly observed in Figure 1b, which shows the propagation of the threshold concentration through the spatial domain at specific times. We observe a significant propagation over the first 10^3^ seconds, which then plateaus to reach a steady state propagation distance of around 0.001 cm at 10^4^ seconds (Figure 1b). Consequently, even if the cell continues secreting for a longer period of time, the threshold concentration could not propagate any farther. We define this maximum propagation distance as the effective signaling distance.

**Table 1:**
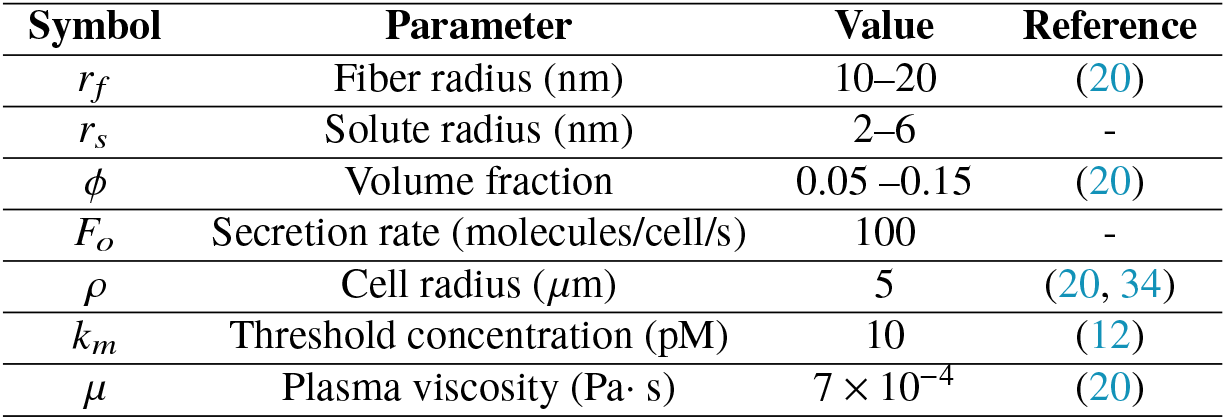
List of parameters used in the model.

**Figure 1:**
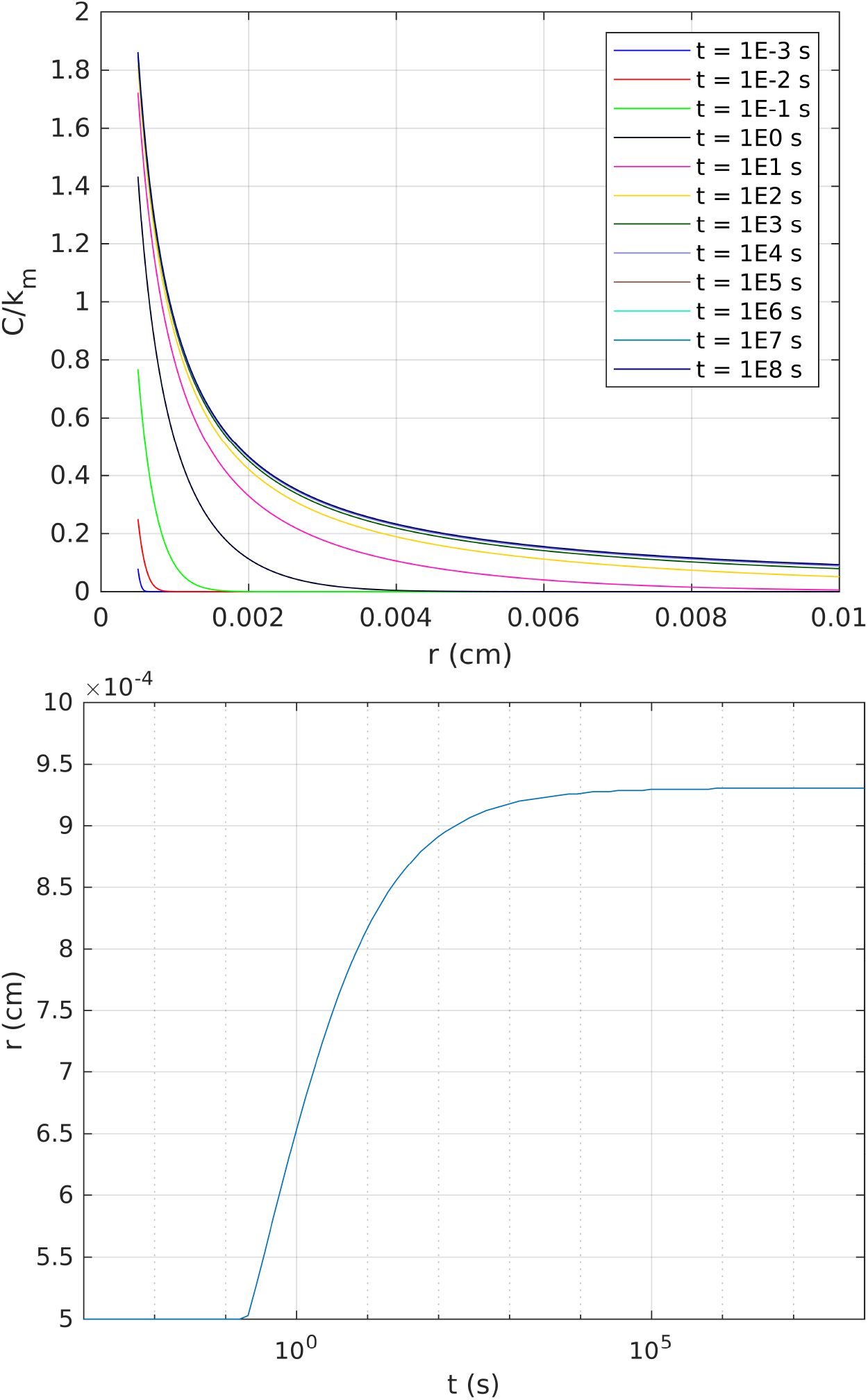
(a) Concentration profile of secreted signaling molecules obtained from analytical solution Eq. 5. Concentration is scaled by the threshold concentration. (b) Propagation distance of the threshold concentration of signaling molecules over time Eq. 6. Base case: *ϕ* of 0.05, *r*_*s*_ of 2 nm, and *r*_*f*_ of 20 nm.

### Collagen Deposition Impact on Diffusivity of Signaling Molecules

The impact of collagen deposition on the effective diffusion coefficient for different sized signaling molecules has been plotted in Figure 2 using physiologically relevant values. A collagen volume fraction of 0.05 represents a healthy matrix, a collagen volume fraction of 0.1 represents a matrix with moderate collagen deposition, and a volume fraction of 0.15 represents a matrix with excessive collagen deposition (20).

**Figure 2:**
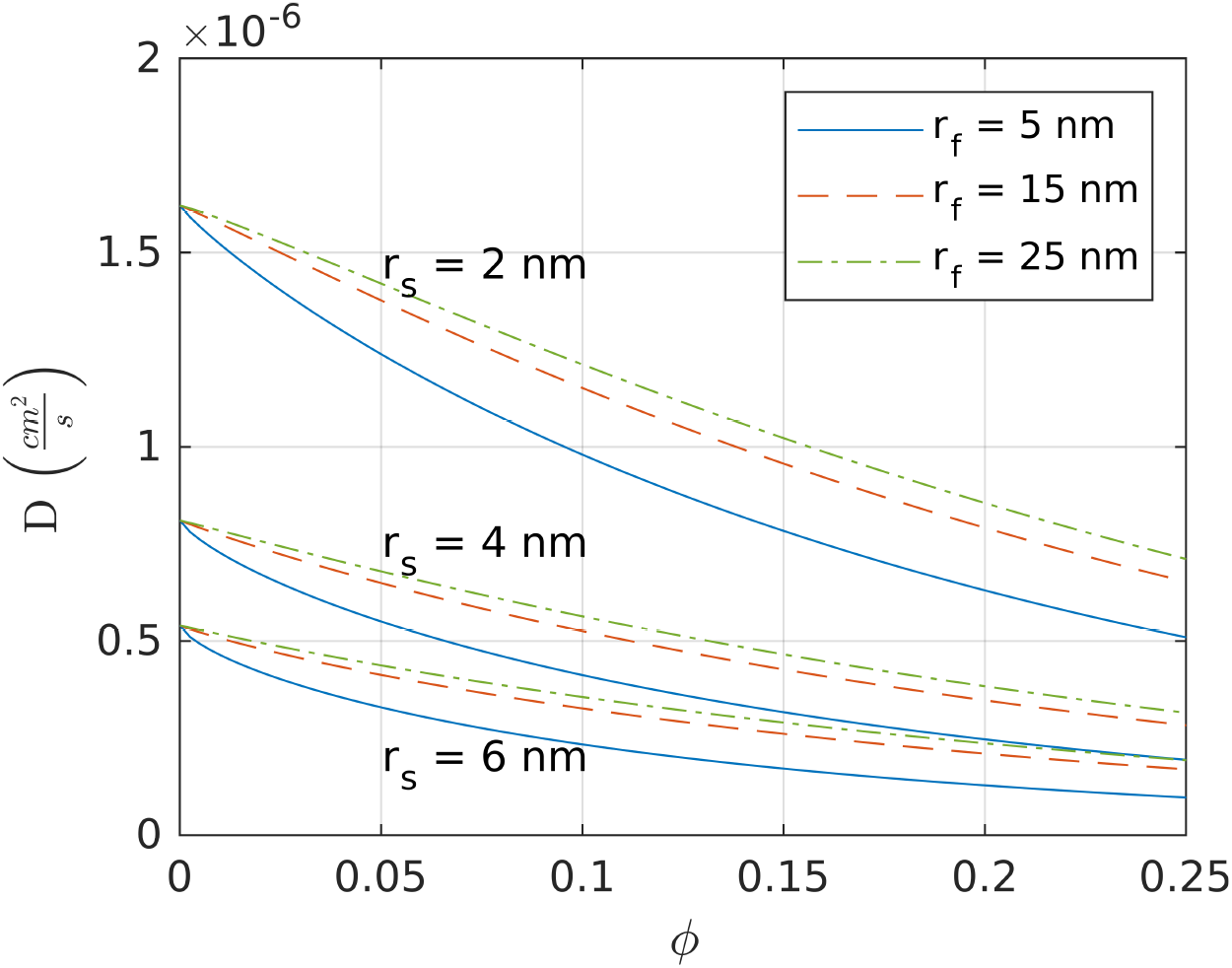
Collagen deposition decreases the diffusivity of signaling molecules. Effective diffusion coefficient *D* as a function of the matrix volume fraction *ϕ*, solute radius *r*_*s*_, and fiber radius *r* _*f*_.

The model shows an exponential decay relationship between the volume fraction and the effective diffusion coefficient that is significant (Figure 2). For pathological changes in fiber radius, the increase in the effective diffusion coefficient is small (Figure 2). Thus, the dependence of the effective diffusion coefficient on the fiber radius is weak relative to the dependence of the effective diffusion coefficient on the volume fraction.

### Signaling Distance Increases with Increased Collagen Deposition

The effective signaling distances for healthy, moderately diseased, and severely diseased matrices for different sized signaling molecules is shown in Figure 3 and represented by the green, yellow, and red curves, respectively. As the matrix becomes diseased due to collagen deposition, the effective signaling distance increases from an initial signaling distance of 2.5 cell radii to approximately 5 cell radii for small molecules and to over 10 cell radii for large signaling molecules (Figure 3). This is surprising, because we had hypothesized that as the matrix becomes diseased, the ability of cells to signal would decrease. However, we see from these results that a diseased matrix actually enhances the ability of a cell to signal rather than diminishing it. Additionally, larger signaling molecules have greater signaling distances than smaller signaling molecules, and the signaling distance of large molecules is further increased in diseased conditions (Figure 3).

**Figure 3:**
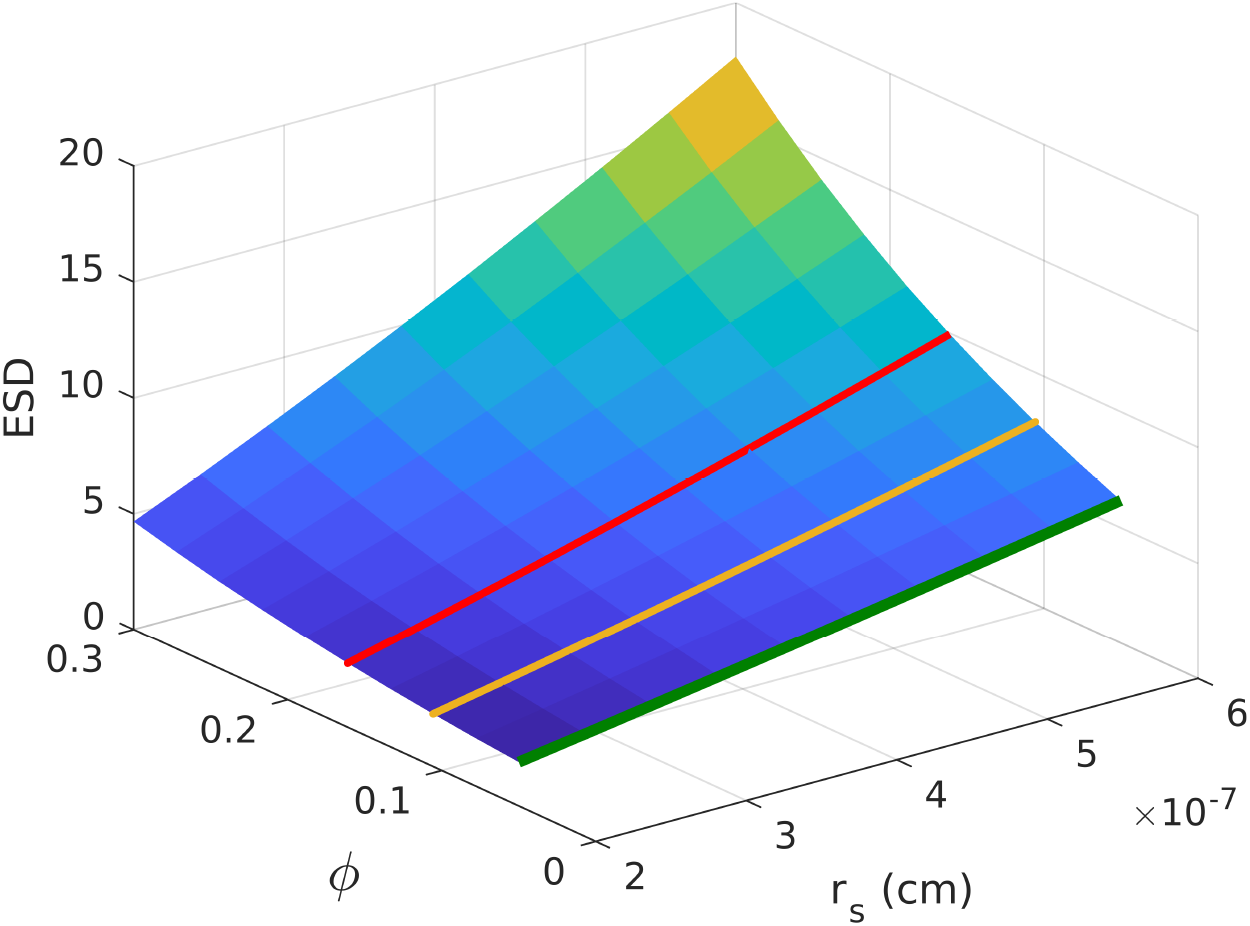
Effective signaling distances for secreted signaling molecules increases as collagen deposition increases. Green, yellow, and red curves represent healthy, moderately diseased, and diseased matrices, respectively. Matrix volume fraction of up to 0.3 is shown to emphasize the nonlinear increasing trend. Results obtained using parameter values listed in Table 1 and *r*_*f*_ of 20 nm.

Compared to previous findings, our predicted values for the effective signaling distance in the healthy case are slightly underestimated. Using experimental and computational methods, Bagnall et al. have estimated that the signaling distance for macrophages is on the order of a few cell diameters (36), a result that has been shown to be the case for Tcells as well (37). Our model also predicts a signaling distance of 2-5 cell radii for the healthy case, but this is an underestimate because we do not incorporate signaling molecule uptake by neighboring cells, which has been taken into account by the other studies. We also assume that no convective transport of signaling molecules occurs which would theoretically increase the signaling distance. Although we cannot specify the extent to which each assumption impacts the effective signaling distance, their opposing influence on effective signaling distance could nullify any individual impact. Regardless, this is a potential area for future work.

Our predictions regarding the changes in effective signaling distance due to collagen deposition in the mesangium, however, cannot be compared to experimental data because such data is not yet available. However, the trend that is observed of an increase in propagation distance due to a decrease in diffusivity is a phenomenon that has been previously described in general (28, 38, 39). Additionally, other modeling studies that consider the uptake of signaling molecules show that the trend is still observed (22). The unexpected increase in signaling distance occurs because the decline in the diffusivity of signaling molecules results in an increased accumulation of signaling molecules at the surface of the cell, which creates a steep gradient leading to a larger propagation distance of the signaling molecules. In our model, this behavior is governed by the boundary condition at the surface of the cell Eq. 2 which states that the concentration gradient at the cell surface is not only a function of the secretion rate but also the diffusion coefficient. Although this modeling strategy has been implemented previously (12, 22), it still needs to be experimentally validated.

### Are There Scenarios Where Signaling Distance Decreases?

Although signaling distance increases with a decrease in diffusivity under both physiological and pathophysiological parameter ranges, there are also cases where signaling distance decreases with a decrease in diffusivity (Figure 4). Concentration profiles of signaling molecules at two diffusion coefficients, representative of matrices with different transport properties and at four separate time points have been plotted (Figure 4). The results show that there are points on the concentration curves, at time 10 s and 100 s, where the higher diffusion coefficient has propagated farther than the concentration curve with the lower diffusion coefficient. However, that is only observed for a short time (Figure 4). As steady state is approached the lower diffusion coefficient propagates further. This behavior is dependent on secretion rate and threshold concentration and has been previously observed theoretically in other biological applications (38, 39).

**Figure 4:**
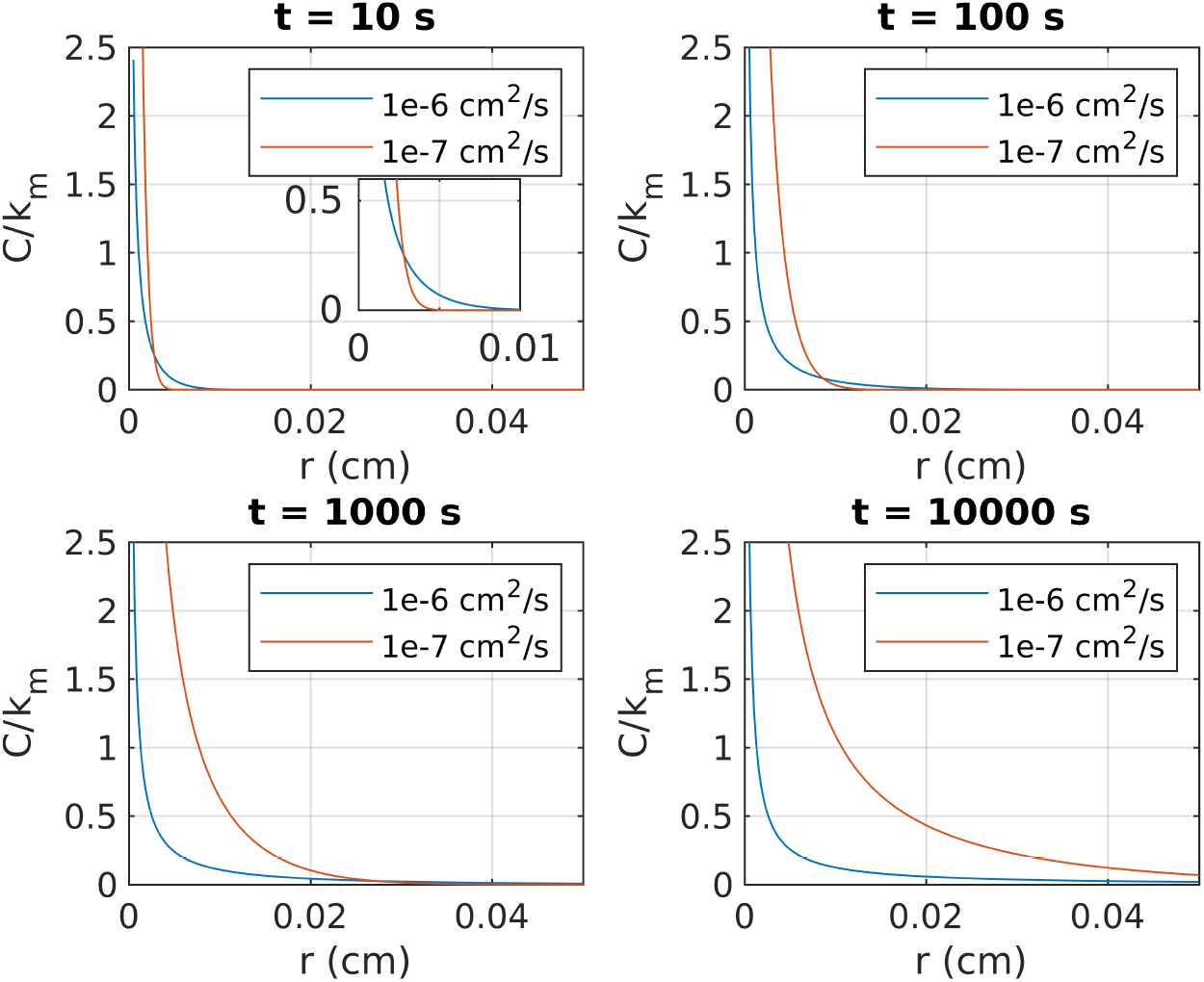
Scenarios where propagation distance of signaling molecules can be reduced for lower diffusivity. *D* = 1 × 10^−6^cm^2^/s represents higher diffusivity, and *D* = 1 × 10^−7^cm^2^/s represents a lower diffusivity.

## CONCLUSION

In this work, we utilized established mathematical models to gain insight into how diabetic kidney damage affects intercellular communication, an important cellular function in health and disease (7–9). More specifically, we elucidated the impact of pathological collagen deposition in the mesangial matrix on intercellular communication. Contrary to our hypothesis, we found that as the level of collagen deposition increases, the signaling range of the mesangial cell increases and that the increase is amplified for large signaling molecules. This finding has implications because an increase in signaling range can result in the propagation of a signal to unintended cells. For example, a profibrotic signal that is normally constrained to immediate neighbors can now propagate to non-neighboring cells to stimulate pathological collagen deposition. This process creates a loop between signaling and collagen deposition that then exacerbates disease condition. In conclusion, our model shows that pathological collagen deposition disrupts the controlled localized signaling of cells, which then contributes to the exacerbation of diabetic kidney damage. This may also hold in other fibrotic diseases.

## ACKNOWLEDGMENTS

This material is based upon work supported by the National Science Foundation under Grant No. 1845117 and resources from Oklahoma State University and the University at Buffalo.

## AUTHOR CONTRIBUTIONS

Haryana Y. Thomas and Ashlee N. Ford Versypt designed the research. Haryana Y. Thomas carried out simulations. Haryana Y. Thomas and Ashlee N. Ford Versypt analyzed the data and wrote the article.

## SUPPLEMENTARY MATERIAL

No supplementary material

